# Microbiome-Dependent Functional Responses to Structurally Distinct Oligosaccharides Revealed by Metaproteomics

**DOI:** 10.1101/2025.06.26.661555

**Authors:** Ailing Zhang, Qing Wu, Janice Mayne, Zhibin Ning, Hongye Qin, Alexander Dewar, Daniel Figeys

**Affiliations:** School of Pharmaceutical Sciences, Faculty of Medicine, University of Ottawa, Ottawa K1H 8M5, Canada; Department of Biochemistry, Microbiology and Immunology, Faculty of Medicine, University of Ottawa, Ottawa K1H 8M5, Canada; Quadram Institute Bioscience, Norwich Research Park, Norwich, Norfolk NR4 7UQ, United Kingdom; University of East Anglia, Norwich, Norfolk NR4 7TJ, United Kingdom

**Author notes:** Corresponding author: Daniel Figeys.

## Abstract

Dietary oligosaccharides are prebiotics that fuel gut microbes, but individual microbiomes may respond differently depending on oligosaccharide structures as well as microbiome composition and function. The extent to which specific gut microbial communities exhibit personalized functional responses to distinct oligosaccharides remains underexplored. We applied a standardized ex vivo microbiome culture, called RapidAIM, coupled with metaproteomics to examine how six structurally diverse oligosaccharides affect the gut microbiota functional response. Our study shows that while human gut microbiomes share some commonalities in utilizing oligosaccharides (e.g. prioritizing dietary fibers over mucin), the fine-scale metabolic and taxonomic responses are highly individualized. Such findings underscore the importance of considering personal microbiome profiles when predicting the outcome of prebiotic interventions. In a broader context, our metaproteomic approach provides a framework for identifying optimal prebiotic choices tailored to individual microbiomes. Ultimately, understanding these personalized responses could inform precision nutrition strategies.

## Introduction

The human gut microbiome, a complex community of microorganisms residing in the gastrointestinal tract, is increasingly recognized as a crucial factor in maintaining host health^1^. It plays vital roles in regulating immune function, aiding in the digestion and absorption of nutrients, synthesizing essential vitamins, and protecting against pathogenic infections^1–3^. The composition and functional capacity of the gut microbiota are influenced by various factors, including diet, genetics, lifestyle, and age^1,4,5^. Disruptions in the balance of this microbial community, known as dysbiosis, have been linked to a range of health issues, such as inflammatory bowel disease, obesity, diabetes, and neurological disorders^6,7^. Consequently, modulating the gut microbiome has emerged as a promising strategy for improving health outcomes.

One such approach is the use of prebiotics, non-digestible food components that selectively stimulate the growth and/or activity of beneficial microorganisms in the gut^8,9^. Among prebiotics, oligosaccharides have attracted considerable attention due to their ability to enhance the proliferation of health-associated bacteria^10^. These compounds are naturally found in breast milk, vegetables, and fruits, and are known to influence gut microbiota primarily through microbial fermentation and utilization.

However, key questions remain unresolved. First, although many studies have investigated the microbial response to oligosaccharide fermentation, they have predominantly focused on compositional shifts in the microbiota^11,12^, in vitro assessments of specific bacterial strains^13^ or functional predictions based on metagenomics^14^. Second, the functional consequences of these interventions, particularly how structurally distinct oligosaccharides are metabolized in vivo, are poorly understood. Third, inter-individual variability in microbial response remains largely unexplored, limiting the ability to design personalized or structure-targeted prebiotic strategies.

Although predictive functional profiling based on metagenomic data has significantly advanced our understanding of microbial communities, it does not accurately predict enzymes levels. Metaproteomics complement these approaches by capturing the expressed proteome, thereby providing more accurate insights into microbial functional responses under specific conditions^15^. Yet, few studies have utilized metaproteomics to simultaneously investigate taxonomic and functional responses of the human gut microbiota to oligosaccharide exposure across individuals^16^. Understanding these responses is critical for optimizing personalized prebiotic strategies. Our study aims to fill this gap by applying metaproteomic analysis to examine how structurally diverse oligosaccharides shape both the taxonomic structure and functional landscape of the human gut microbiome, with a particular focus on inter-individual differences.

## Results

### Individual Differences in Gut Microbiota Composition Among Three Females

Briefly, six structurally distinct oligosaccharides were used in this study: *1,5-α-L-arabinotetraose* (ARA), *1-kestose* (KES), *D-(+)-raffinose pentahydrate* (RAF), *isomaltotriose* (ISO), *D-cellotriose* (CEL), and *1,4-β-D-xylotriose* (XYL3). The effects of these prebiotics, along with PBS (as vehicle control and baseline reference), were evaluated on microbiomes derived from three healthy adult donors (F1, F2, and F3) using the ex vivo RapidAIM^17^ assay in triplicate. After 18 hours of prebiotic exposure, the microbiomes were lysed and the proteins processed for metaproteomic analysis (Figure 1a). A total of 1,365,863 peptides, and 21,068 quantified proteins were observed across 63 samples. As well, 792 species from 385 genera were identified (Figure 1b).

**Figure 1.**
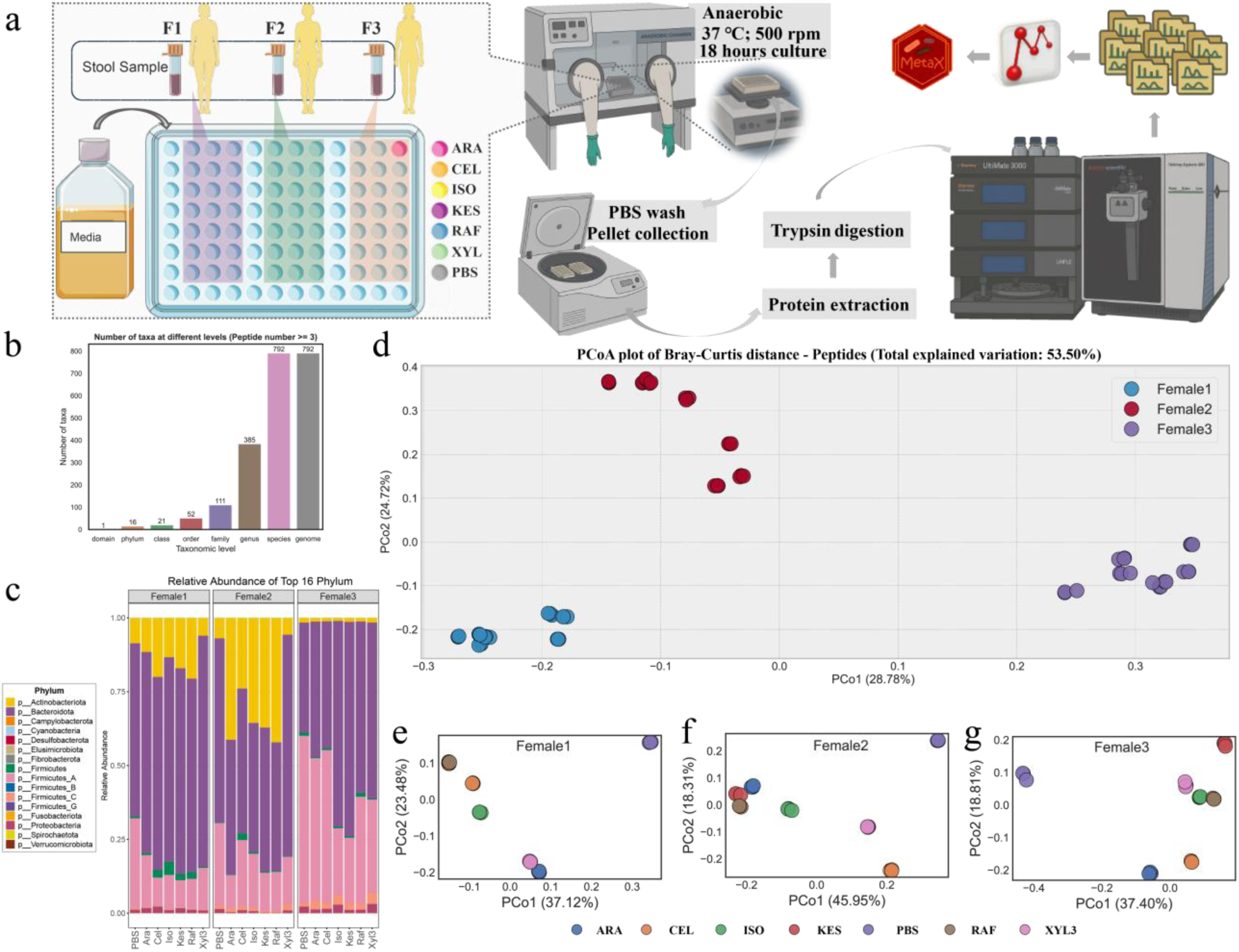
Overview of experimental workflow and taxonomic diversity across treatments. (a) Summary of workflow. (b)Total number of identified taxa across different taxonomic levels. (c) Relative abundance at the phylum level under various treatments. (d) Peptide-based distributional distances among all samples. (e–g) Peptide-based distributional distances for F1, F2, and F3, respectively.

The baselines functional composition of the three microbiomes are different and this difference remains after treatments with different oligosaccharides (Figure 1d). When analyzed individually, the six oligosaccharide-treated samples tended to cluster more closely together in the PCoA distance, while the PBS control sample was consistently positioned further apart (Figure 1e, f, g). This indicates that oligosaccharide treatments induced a shift in gut microbial composition relative to the baseline. However, the clustering patterns among the six treatments varied across individuals.

As expected, substantial inter-individual differences in gut microbiome functional composition were observed among the three adult females. Notably, Actinobacteriota exhibited much lower relative abundance in F3 compared to F1 and F2, both of whom showed relatively high levels that further increased upon treatment with all oligosaccharides except XYL3 (Figure 1c), with F2 showing a greater increase than F1. At baseline, Bacteroidota was more abundant in F1 and F2 than in F3. Following oligosaccharide treatments, its abundance decreased in F2 (except under XYL3) but increased in F1 and F3. Firmicutes_A was the most dominant phylum overall, yet its relative abundance decreased in all individuals after treatments (Figure 1c).

### Oligosaccharide-Induced Activation of Biosynthetic Functions and Suppression of Stress-Responsive Pathways

In our experiment we specifically selected oligosaccharides with distinct glycan compositions and structures, each representative of different oligosaccharide families. Given this diversity, we first examined whether functional expression was: (i) commonly affected by the oligosaccharides across all individuals, (ii) specifically influenced by each type of oligosaccharide, and/or (iii) uniquely modulated in an individual-specific manner. We observed consistent expression patterns across different treatments within the same individual (in at least three treatment conditions) (Supplementary Data S1). Differentially expressed KEGG modules were identified and summarized as follows: 24 functions were upregulated and 26 were downregulated in F1; 29 functions were upregulated in both F2 and F3. Notably, F2 exhibited the highest number of downregulated functions (32), while F3 shared the same set of downregulated functions as F1. Three KEGG modules were consistently upregulated across all three individuals following oligosaccharide treatments: M00028 (ornithine biosynthesis), M00093 (phosphatidylethanolamine biosynthesis), and M00432 (leucine biosynthesis) (Supplementary Data S1).

Conversely, eight functional modules were commonly downregulated across most treatments in all three individuals (Supplementary Data S1). These downregulated functions were primarily involved in complex carbohydrate degradation, energy-saving or stress-responsive metabolism, and aromatic compound degradation.

### Functional Responses Reflecting Donor-Specificity to Oligosaccharide Treatments

Despite the observation of several shared functional changes, we also identified individual-specific functional responses of the gut microbiota to different oligosaccharide treatments (Figure 2a). In F1, a subset of consistently regulated functional modules—present across multiple oligosaccharide treatments (at least three treatments)—was characterized by the upregulation of amino acid degradation pathways, including the GABA shunt and branched-chain amino acid catabolism, as well as sphingolipid degradation and pantothenate biosynthesis. Concurrently, modules involved in the metabolism of complex carbohydrates, such as the uronate pathway, and non-phosphorylative Entner-Doudoroff (ED) pathway were downregulated (Figure 2b). F2 displayed a subset of upregulated modules across multiple treatments that highlighted increased biosynthetic capacity, including variants of the pentose phosphate pathway (PPP), several amino acid biosynthesis routes (e.g., methionine, threonine, valine), nucleotide biosynthesis, triglyceride formation, and vitamin/cofactor pathways (e.g., folate, menaquinone, ascorbate). Simultaneously, reductions in purine degradation, polyamine biosynthesis, LPS core synthesis, and methanogenesis were noted (Figure 2c). In F3, several consistently upregulated pathways across multiple treatments were related to fermentation-associated metabolism (e.g., modules involving methanol and methylamine utilization), KDO₂-lipid A biosynthesis, biotin production, and fatty acid synthesis. At the same time, pathways involved in central carbon metabolism—including the PPP, TCA cycle entry steps, and biosynthesis of proline and GABA—were downregulated (Figure 2d).

**Figure 2.**
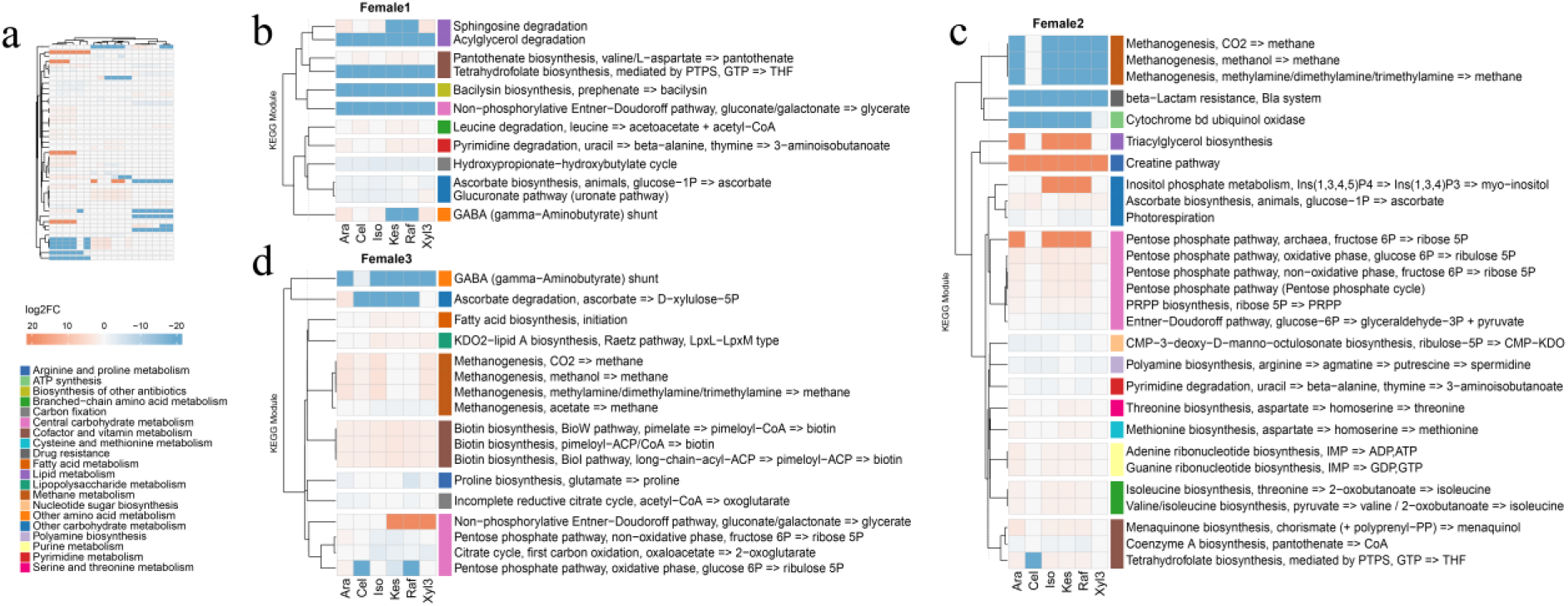
KEGG-based functional profiling reveals individual-specific yet stable treatment responses. (a) Schematic showing consistent functional expression within individuals across treatments. (b–d) Functional module profiles for each individual, highlighting intra-individual consistency and inter-individual divergence. Colored blocks on the right indicate corresponding Level 3 KEGG functional categories.

Interestingly, some oligosaccharide treatments led to responses that were different from most of the others within the same individual. In F1, the GABA shunt and sphingolipid degradation were downregulated in response to KES and RAF, but upregulated with ARA, ISO, and XYL3. In F2, only CEL caused a reduction in folate biosynthesis, whereas other oligosaccharides increased it. This may suggest that CEL was utilized by microbial groups that do not contribute to folate production. In F3, the PPP and ascorbate degradation were upregulated only under ARA treatment, likely due to its unique structure and direct metabolic routing.

### Oligosaccharide-Driven Upregulation of initial cleaving CAZymes and Suppression of Mucin degradation

At a functional level, microbial responses to oligosaccharides can be further understood by examining pathways involved in carbohydrate metabolism. The first enzymatic step when microbiomes are exposed to oligosaccharides is to cleave the glycosidic bonds in the polymer leading to monomers. Some bacteria are capable of processing oligosaccharides through the action of a subset of enzymes classified as carbohydrate-active enzymes (CAZymes)^18^. To investigate the gut microbiota’s response to various oligosaccharides, we identified CAZymes whose abundance changed consistently across all samples after exposure to each oligosaccharide. We then mapped and visualized the distribution of these CAZymes responsible for the initial cleavage of glycosidic bonds (Figure 3a, b). As expected, these CAZymes showed substrate-specific induction based on the structure and composition of the applied oligosaccharides, facilitating their breakdown into smaller units that can be further metabolized or structurally utilized by the microbiota, including but not limited to use as carbon sources. Briefly (Figure 3a, b, c), ARA induced the upregulation of arabinosidase^19^ (GH51/GH43_4), KES upregulated fructosidase (GH32), RAF upregulated alpha−galactosidase^20^ (GH36) and GH32, ISO upregulated alpha−glucosidase (GH31) and oligo−1,6−glucosidase^21^ (GH13_31), CEL upregulated cellulase (GH5_2) and beta-glucosidase (GH3), and XYL3 upregulated xylanase (GH8) and xylosidase^22^ (GH43_12 and GH43_1). These patterns confirm that the type of oligosaccharide determines the specific CAZymes being upregulated. Notably, this substrate-specific response was observed consistently across all three individuals, indicating a shared functional adaptation of their microbiota (Figure 3a, b).

**Figure 3.**
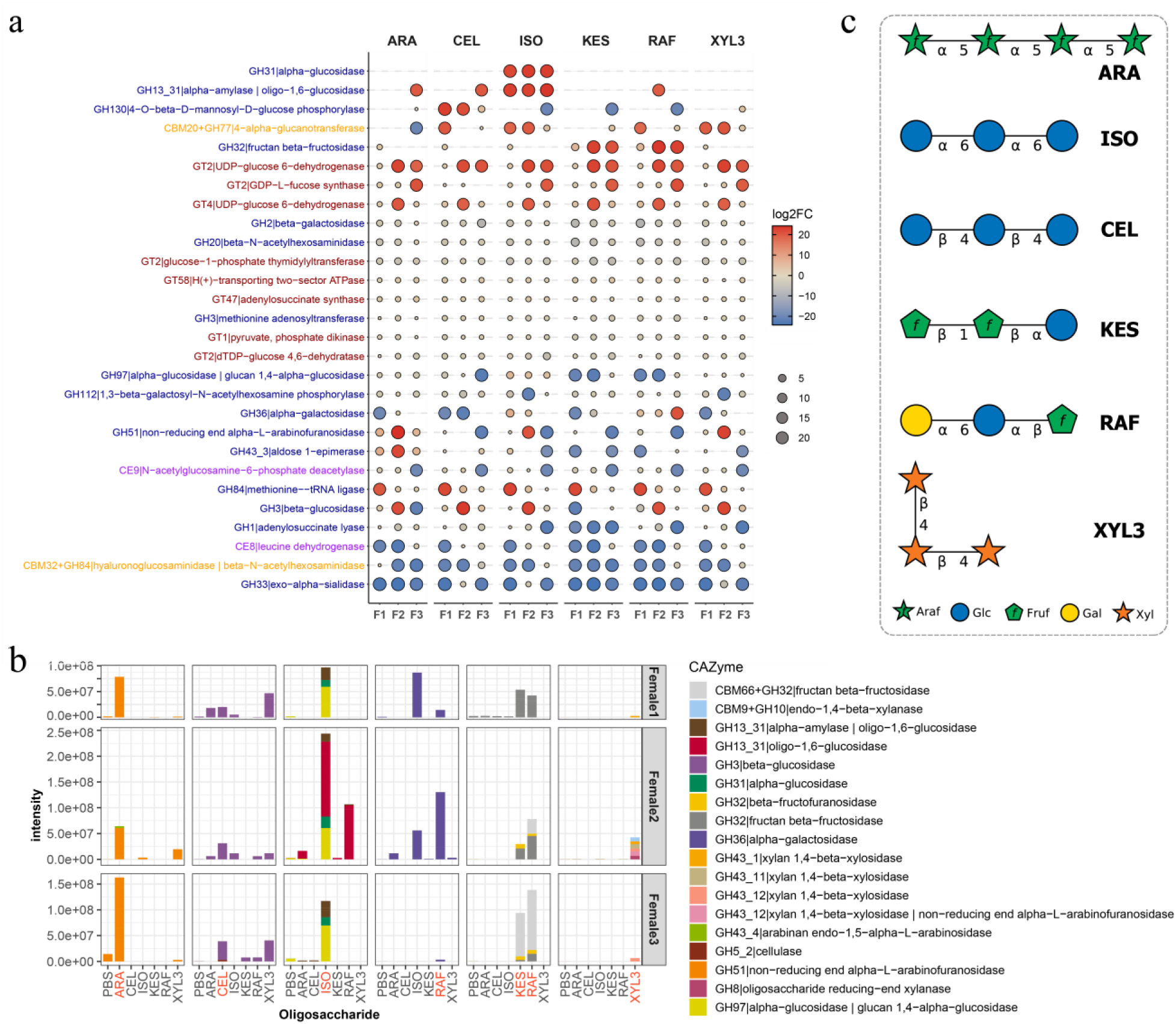
Consistent induction and abundance of oligosaccharide-degrading CAZymes across treatments. (a) Significantly expressed oligosaccharide-cleaving CAZymes showed consistent expression patterns under specific treatments across all three individuals. (b) Intensity of oligosaccharide-cleaving CAZymes in each individual under different treatments. The treatment being applied in each condition is marked in red. (c) Biologic description of oligosaccharides from PubChem. ARA: CID 72734308; ISO: CID 439668; CEL: CID 5287993; KES: CID 440080; CID 439242; XYL3: CID 10201852^24^.

A notable common feature among all treatments was the down-regulation of mucus-degrading enzymes (e.g., GH33, GH84) ^23^ (Figure 3a). This suggests that regardless of the type of exogenous oligosaccharide, if it can be utilized by the microbiota to provide energy and substrates, then microbial dependence on host mucus is reduced. Additionally, all six oligosaccharides reduced the activity of enzymes such as N-acetylhexosaminidase (Figure 3a).

Although similar expression patterns of these initial CAZymes were observed across the three individuals, donor-specific differences were also evident. For example, three fructan-related enzymes were detected following KES/RAF treatment. However, fructan β-fructosidase (CBM66+GH32/GH32) was present in F2 and F3, while F1 only expressed β-fructofuranosidase. In addition, six different CAZymes associated with XYL3 degradation responded to treatment in F2, whereas only one was detected in F1 and F3, respectively (Figure 3b). Beyond these substrate-specific responses, some unexpected expression patterns were also observed. For instance, GH32 was upregulated under all treatments in F1, despite its known specificity for fructans such as KES and RAF (Figure 3a, c).

### Cross-Individual Functional Redundancy in CAZyme Expression

We linked the CAZymes to their corresponding taxa using MetaX^25^, allowing us to identify the potential microbial contributors to primary oligosaccharide degradation. We found those initial CAZymes mainly produced from six bacteria families (Bifidobacteriaceae, Lachnospiraceae, Bifidobacteriaceae, Ruminococcaceae, Muribaculaceae, and Rikenellaceae) containing 15 genus (*Bariatricus*, *Blautia_A*, *Bacteroides*, *Anaerosacchariphilus*, *Mediterraneibacter*, *Prevotella*, *Phocaeicola*, *Bifidobacterium*, *Gemmiger*, *Agathobacter*, *CAG-873*, *Dorea_A*, *Alistipe*s, *Anaerostipes*, *Faecalibacterium*) and 33 species (Figure 4b).

**Figure 4.**
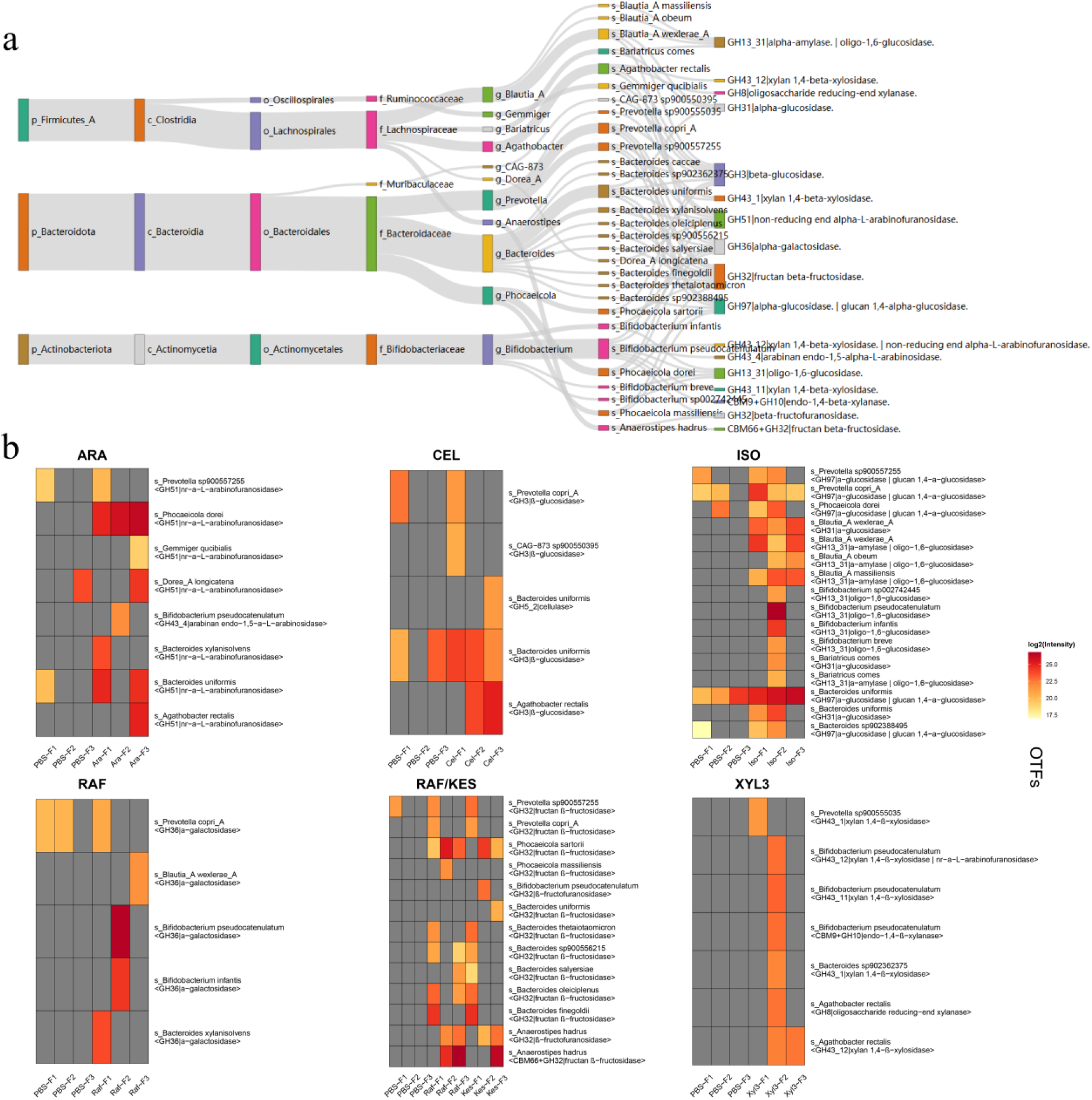
Visualization of Operational Taxa-Function (OTF) expression profiles and their taxonomic origins. (a) Sankey diagram linking CAZymes to their corresponding taxa. (b) Heatmap of OTFs (Operational Taxa-Functions) representing oligosaccharide-cleaving CAZymes across different treatments and individuals. Rows indicate taxa-function pairs (OTFs), and columns represent specific treatment conditions per individual. Color intensity reflects the log2(expression level) of each OTF.

We observed that under ISO treatment, alpha-glucosidase activity (GH31) was primarily attributed to *Bacteroides uniformis*, *Blautia_A wexlerae_A*, and *Bariatricus comes*. However, only two of these species were responsible for GH31 activity in F1, and only one in F3. In contrast, GH13_31 (annotated as alpha-amylase or oligo-1,6-glucosidase) was produced by eight species in F2, predominantly from the *Blautia_A* and *Bifidobacterium* genera, whereas only two and three *Blautia_A* species were detected with this CAZyme activity in F1 and F3, respectively. A similar pattern was observed in response to xylotriose treatment. In F1 and F3, only one species was associated with xylotriose-related CAZyme activity—*Prevotella sp900555035* (GH43_1) and *Agathobacter rectalis* (GH43_12), respectively. In contrast, six species in F2, belonging to the *Bacteroides*, *Bifidobacterium*, and *Agathobacter* genera, were involved in the degradation of xylotriose. Notably, *Bifidobacterium pseudocatenulatum* in F2 expressed three different xylotriose-targeting CAZymes (Figure 4b).

These patterns were also observed across other CAZymes, with *Bacteroides*, *Bifidobacterium*, and *Prevotella* playing central roles in carbohydrate metabolism. Importantly, F2’s microbiota exhibited notably higher *Bifidobacterium*-associated enzymatic activity across multiple targets, highlighting inter-individual variation in oligosaccharide metabolism (Figure 4b).

### Microbial Context Determines Individualized Metabolic Processing of Oligosaccharides

We examined the changes in the relative abundance of genera capable of producing specific CAZymes under a particular treatment and observed that the expression of these initial CAZymes was generally accompanied by an increase in the corresponding genera with a few exceptions. Notably, the taxonomic shifts displayed strong individual specificity across different treatments, consistent with the individual-dependent patterns we previously observed in overall functional changes. We observed that in F2, *Bifidobacterium* became dominant in relative abundance following most treatments, accompanied by a more pronounced shift in microbial composition compared to her baseline. This shift was notably greater than those observed in F1 and F3. Specifically, the dominant genus transitioned from *Bacteroides* at baseline to *Bifidobacterium* after treatment (Figure 5a).

**Figure 5.**
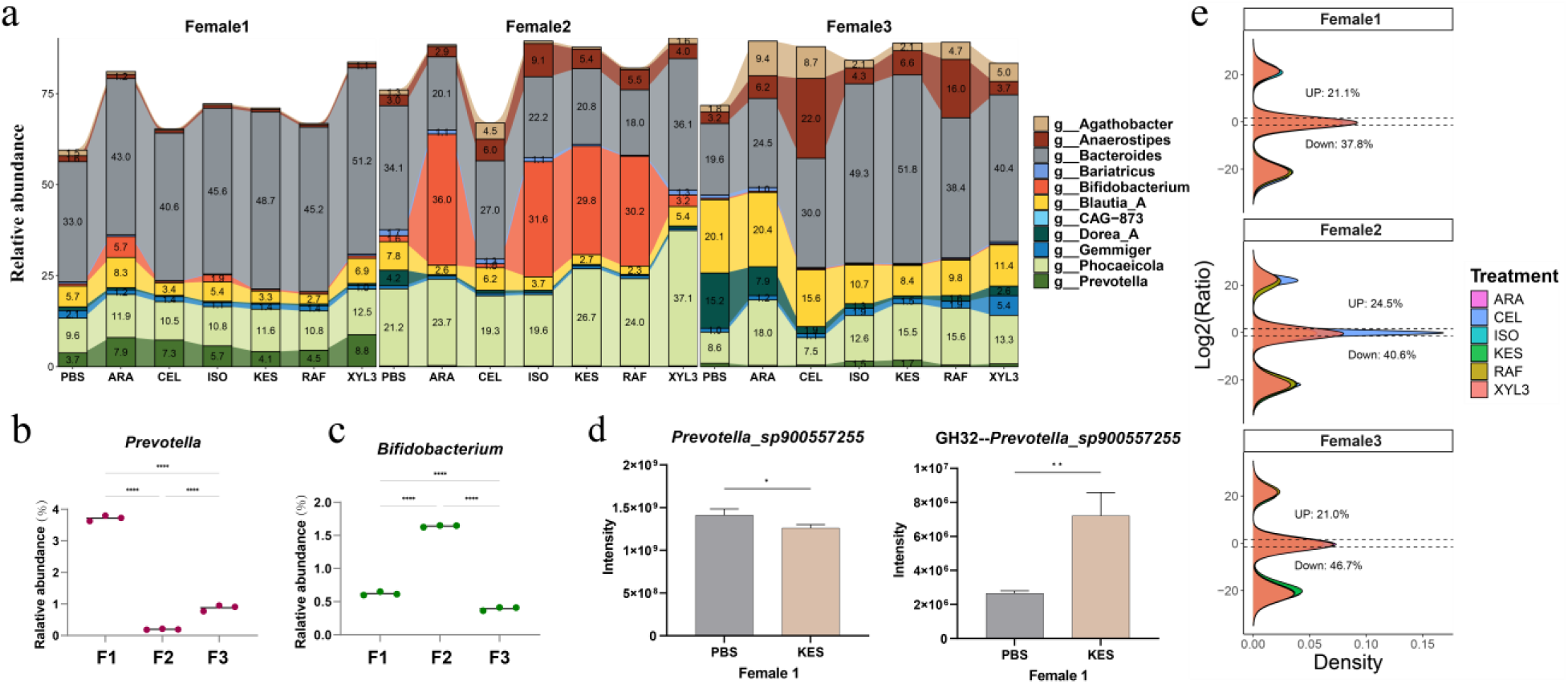
Taxonomic-functional dynamics of oligosaccharide-responding gut microbes. (a) Genus-level taxonomic composition of gut microbiota in each individual across various oligosaccharide treatments. Stacked area plots show the relative abundance (%) of CAZymes produced genera under control (PBS) and oligosaccharide-treated conditions (ARA, CEL, ISO, RAF, KES, XYL3). (b) Baseline relative abundance of *Prevotella*. (c) Baseline relative abundance of *Bifidobacterium*. (d) Intensity of *Prevotella_sp900557255* and GH32 it produced in Female 1 under KES treatment and PBS. (e) Global overview of the ratio of functional change to the corresponding taxonomic change across three individuals. Horizontal dashed lines indicate thresholds of log₂(Ratio) = ±1.5.

### Baseline Relative Abundance Determines CAZyme Upregulation, Mediating Genus-Level Abundance Increase

As we further explored the causes of functional redundancy, we observed that CAZyme production is broadly associated with the relative abundance of different microbial genera in each individual. Specifically, under various oligosaccharide treatments, the genera responsible for producing key enzymes often had relatively high baseline abundances (Figure 5a).

For instance, in the control group for ARA treatment, we found that the relative abundance of *Bacteroides* and *Phocaeicola* was high across all three individuals. Correspondingly, we observed an upregulation of alpha-L-arabinofuranosidase in these two genera (with the exception of *Bacteroides* in F2), and an increase in their relative abundances post-treatment. By contrast, the baseline relative abundance of *Prevotella* was 3.7%, 0.20%, and 0.88% in the three individuals, respectively, with a markedly higher abundance observed in F1 compared to the other two (Figure 5b). GH51 production from *Prevotella* was only detected in F1, where an increase in *Prevotella*’s relative abundance was also observed only in F1 after ARA treatment (Figure 5a). Moreover, *Agathobacter* and *Dorea_A* exclusively produced alpha-L-arabinofuranosidase in F3. In this individual, *Agathobacter* exhibited a higher baseline abundance and showed a further increase after treatment, whereas *Dorea_A* decreased. Interestingly, a reduction in *Dorea_A* was also observed in the other two donors following treatment with ARA, and even across all oligosaccharide treatments (Figure 5a). This may suggest that the decline of *Dorea_A* is a general response to these oligosaccharides, potentially linked to substrate competition, selective pressure, or ecological displacement within the microbial community. *Bifidobacterium* had significant higher baseline relative abundances in F2 (0.62%, 1.64%, and 0.40% across the three individuals, respectively) (Figure 5c), and we only detected endo-1,5-alpha-L-arabinosidase expression in F2 (Figure 4b). Despite the generally low abundance of this genus, we observed a sharp increase in *Bifidobacterium* in F2 post-treatment (Figure 5a).

However, in F2, we did not observe the expected increase in *Bacteroides* abundance despite the high relative abundance shown in baseline (Figure 5a). One possible explanation involves the notably higher baseline abundance of *Phocaeicola*, particularly *Phocaeicola dorei*, which showed the highest expression level of alpha-L-arabinofuranosidase across all individuals, suggesting a strong capacity for arabinose release. In addition, we detected exclusive expression of endo-1,5-alpha-L-arabinanase (EC 3.2.1.99) in F2 (Figure 4b). This enzyme cleaves the arabinofuranosyl backbone of arabinan into arabino-oligosaccharides, which are further hydrolyzed into L-arabinose by alpha-L-arabinofuranosidase (EC 3.2.1.55)^26^. Such a combination of enzyme activity may promote more efficient degradation of ARA, potentially facilitating rapid utilization by *Bifidobacterium*. The observed increase in *Bifidobacterium* abundance in F2, from 1.64% to 36.01% (Figure 5a), reflect a competitive advantage in utilizing released sugars.

Except for the ARA treatment, a sharp increase in *Bifidobacterium* abundance was observed under KES, RAF and ISO treatment. To further investigate the underlying mechanisms, we looked at the enzymes the different bacteria produced in response to the same oligosaccharide. Although functional redundancy was apparent, where distinct taxa produced enzymes with similar activities, differences in enzyme specificity and expression patterns were also evident. For instance, in response to ISO, *Bifidobacterium* predominantly expressed oligo-1,6-glucosidase (GH13_31), an enzyme with high specificity for α-1,6-glucosidic linkages, the structural basis of ISO. In contrast, Bacteroidaceae and Lachnospiraceae responded to ISO by expressing alpha-glucosidase (EC 3.2.1.20), an enzyme with broader substrate specificity and less efficiency toward α-1,6-glucosidic bonds^27^ (Figure 4b).

Under KES treatment, *Bacteroides* and *Phocaeicola* mainly produced fructan beta-fructosidase (EC 3.2.1.80), whereas Bifidobacterium expressed beta-fructofuranosidase (EC 3.2.1.26). Although both enzymes belong to the GH32 family, their substrate preferences diverge: fructan beta-fructosidase is generally more efficient at degrading long-chain fructans (e.g., inulin), whereas beta-fructofuranosidase is more active on short-chain FOS with β-2,1 linkages^28^. We also observed that *Anaerostipes* expressed beta-fructofuranosidase under KES treatment in both F2 and F3 samples. However, in F2, the expression level of this enzyme by *Bifidobacterium* far exceeded that of *Anaerostipes*. A similar pattern was observed under RAF and XYL3 treatment, where multiple taxa encoded functionally similar enzymes, yet the dominant contributors differed in abundance and taxonomic affiliation (Figures 4b, 5a).

### Metaproteomic Evidence of Flexible Enzyme Expression Across Oligosaccharide Treatments

Some species were found to produce different enzymes in response to distinct oligosaccharide treatments, especially *Bifidobacterium pseudocatenulatum*, which positively responded to all oligosaccharides in F2 (Figure 4b). *Bacteroides uniformis* produced alpha-L-arabinofuranosidase in F1 but produced fructan beta-fructosidase in F3. Likewise, *Agathobacter rectalis* contributed alpha-L-arabinofuranosidase in F3 and 1,4-beta-xylosidase in Females 2 and 3 (Figure 4b). These results suggest that these species play a crucial role in oligosaccharide metabolism due to their flexible enzyme production in response to different treatments.

Our results also indicate that changes in functional activity are not always directly coupled with changes in taxonomic abundance. *Dorea_A*, as we discussed, that despite expressing alpha-L-arabinofuranosidase in F3, decreased in intensity following treatment (Figures 4b, 5a). Besides, *Prevotella* under KES treatment exhibited a significant reduction in abundance compared to PBS, while its associated enzyme GH32 was strongly upregulated (Figure 5d). To further illustrate the relationship between taxa based functional change and taxa change in our metaproteomics data, we compared the intensity change of taxa and their corresponding function proteins, and a global change ratio was shown in Figure 5e. ∼ 21% to 25% (log₂ ratio > 1.5) of proteins were significantly higher than the changes of taxa across three individual microbiomes while 37% to 47% showed opposite trend (log₂ ratio < −1.5) between function change and taxa change. This means most of the protein changes were not explained by taxa changes.

### Strong Capacity of Bacteroidaceae for Xylotriose Degradation and Utilization

Although we found different kinds of XYL3 related CAZymes expressed in F2, we did not observe a significant increase in the relative abundance of *Bifidobacterium* like other treatments. To further investigate the potential reasons, we applied weighted gene co-expression network analysis (WGCNA)^29^, grouping proteins into distinct co-expression modules based on expression patterns, with certain modules showing strong correlations with specific oligosaccharide treatments (Figure 6a). Among the results, we found significant enrichment of CAZymes responsible for cleaving glycosidic bonds, highlighting the tight associations between specific protein modules and oligosaccharide treatments. In addition, we identified xylose isomerase activity consistently across all three individuals, indicating the bacterial ability to utilize D-xylose in energy production (K01805: xylA; xylose isomerase [EC:5.3.1.5]). For xylose isomerase, we identified upregulation across eight genera. Notably, *Bacteroides* and *Phocaeicola* exhibited high levels of this enzyme across all three individuals, indicating a consistent role in D-xylose metabolism (Figure 6b). Additionally, *Bifidobacterium* demonstrated upregulated expression of xylose isomerase in both F1and F2. We next identified a xylose transporter, xylE (K08138; MFS transporter, SP family, xylose⁺ symporter), belonging to the major facilitator superfamily (MFS) (Figure 6c). Interestingly, this transporter showed higher abundance in F1and F3 compared to F2. Further examination revealed that the xylE transporters were derived from three species within the genera *Bacteroides* and *Phocaeicola* (*Bacteroides uniformis*, *Bacteroides xylanisolvens*, and *Phocaeicola dorei*). Notably, xylE from *Phocaeicola* was exclusively found in F2, while xylE from *Bacteroides uniformis* was detected in F1and F3. This finding aligns with the observed changes in the relative abundance of these two genera under XYL3 treatment (Figure 6d). This may potentially explain why, under the XYL3 treatment, the relative abundance of *Bacteroides* did not decrease in F2, further highlighting the strong capacity of Bacteroidaceae for xylotriose degradation and utilization.

**Figure 6.**
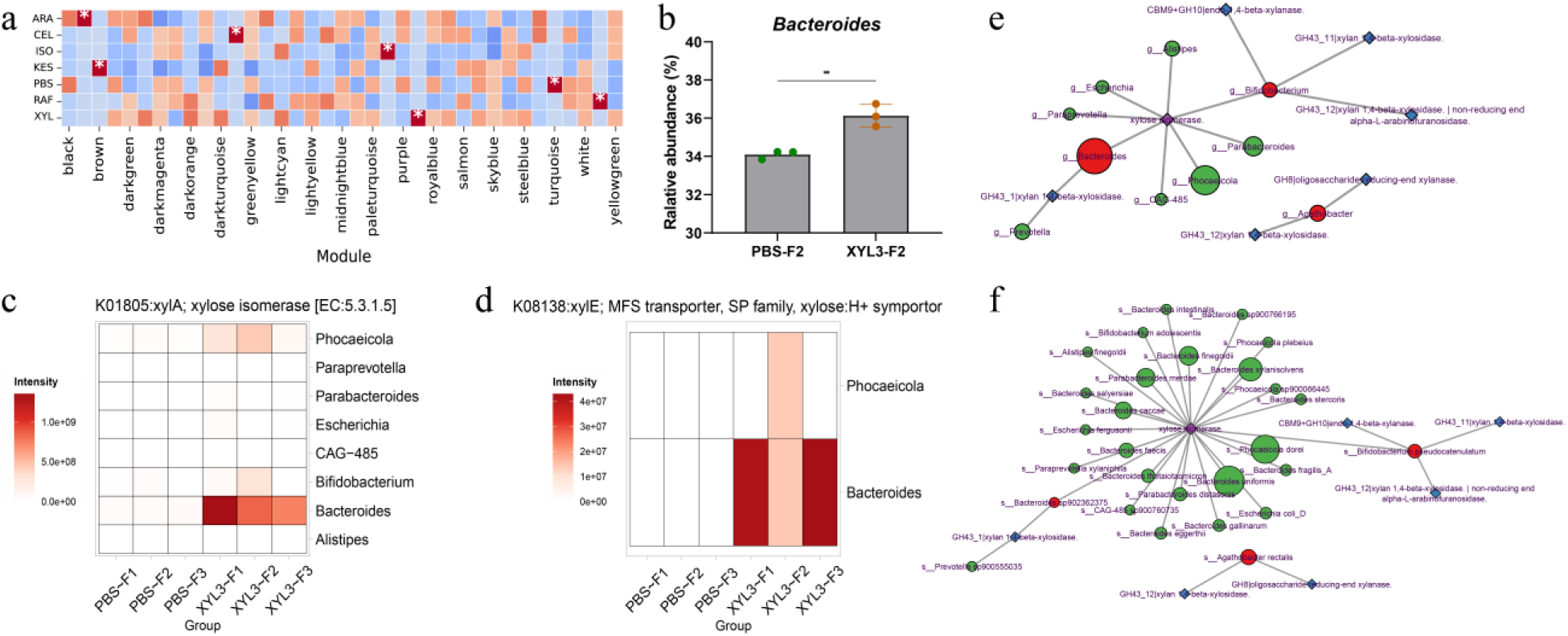
Functional and taxonomic coordination in microbial response to xylotriose. (a) Schematic representation of WGCNA results (F3). (b) Relative abundance of *Bacteroides* in F2 under PBS and xylotriose treatments. Asterisks indicate significant differences between the two groups. (c) Expression levels of xylA under control and xylotriose treatments. (d) Expression levels of xylE under control and xylotriose treatments. (e) A taxa-function network at the genus level illustrating the producers of both primary (blue diamonds) and secondary (purple diamonds) enzymes involved in xylotriose metabolism after treatment. Genera capable of producing primary CAZymes are labeled in red, while those producing only secondary enzymes are labeled in green. (f) The same network as in (e) but presented at the species level.

### Cross-feeding via downstream Carbohydrate Metabolism

Our earlier results indicated that the initial cleavage of oligosaccharides is carried out by a subset of microbial taxa through the expression of specific CAZymes. This raised the question of whether the downstream utilization of the resulting monosaccharides (or terminal sugar residues) is also restricted to the same taxa or is leading to cross-feeding^30^. To address this, we examined the taxa responsible for expressing secondary enzymes involved in monosaccharide metabolism. Our analysis revealed that, across several treatments, cross-feeding was occurring with community members responsible for downstream monosaccharide metabolism extended beyond the initial oligosaccharide degraders.

For example, under XYL3 treatment, we identified three species from three different genera as the primary producers of xylan-degrading CAZymes. In contrast, the enzyme xylose isomerase, which converts xylose into xylulose for further metabolism, was detected in 24 species spanning six additional genera that were not involved in the initial degradation step (Figure 6e, f). This finding suggests that cross-feeding is occurring. While a few taxa initiate the breakdown by producing primary CAZymes, the resulting monosaccharides or residues can be further utilized by a wider range of species. This indicates a functional specialization within the microbial community, where more members are involved in later stages of carbohydrate metabolism than in the initial degradation step. Notably, *Agathobacter*, one of the primary xylan degraders, did not express xylose isomerase.

## Discussion

In this study, we investigated how three individual gut microbiomes functionally respond to six structurally distinct oligosaccharides using a metaproteomic approach. Bacteroidota, Firmicutes_A and Actinobacteriota are often reported to dominate the gut microbiota, which are known for their roles in gut homeostasis and commonly used as probiotics due to their health-promoting effects^31,32^. In our study, however, these phylum-level compositions and changes likely reflect individual-specific adaptations of the gut microbiota to oligosaccharide treatments, highlighting the idea that gut microbiota responses to prebiotics are highly personalized. Consistent with these taxonomic differences, the functional module-level responses also exhibited substantial inter-individual variability. For example, although all three microbiomes exhibited changes in carbohydrate metabolism following oligosaccharide treatment, the preferred metabolic routes involved—such as the ED pathway, PPP, or TCA cycle—varied between individuals. A study in a rat model showed that intake of slow-digesting carbohydrates can concurrently upregulate all three pathways^33^, suggesting that carbohydrate structure broadly modulates central carbon metabolism. Our results indicate that while carbon utilization was commonly affected, the metabolic routes engaged were highly individual-specific. Similarly, vitamin biosynthesis pathways, including those for B5, B7, B9, and K2, exhibited subject-specific regulation patterns. Microbial synthesis of these vitamins has been reported in both animals and humans^34^, and our findings suggest that oligosaccharide exposure may influence microbial vitamin production in a host-specific manner. Although each individual displayed some consistent functional changes across different oligosaccharide treatments, there were also clear treatment-specific differences. Some opposite trends in certain treatments (e.g. only CEL caused a reduction in folate biosynthesis, whereas other oligosaccharides increased it in F2), show that individual responses are still influenced by the structure of each oligosaccharide. This suggests that even in a personalized microbiota, the chemical features of each oligosaccharide play a key role in shaping the functional outcome.

Despite these inter- and intra-individual differences, we identified a set of functional modules that were consistently regulated across all individuals and oligosaccharide treatments. Modules associated with amino acid biosynthesis and membrane biogenesis were commonly upregulated, suggesting a shared shift toward growth-associated metabolic activity. Conversely, functional modules involved in complex carbohydrate degradation and energy conservation, were generally downregulated. This consistent pattern implies that, regardless of individual microbiome composition or oligosaccharide structure, the gut microbiota collectively adopted a metabolic strategy favoring biomass accumulation and reduced reliance on stress-related pathways under oligosaccharide-rich conditions.

Among the functional categories involved in dietary carbohydrate utilization, CAZymes play a central role in initiating the breakdown of complex oligosaccharides^18^. We found that under all six oligosaccharides treatments, mucus-degrading related CAZymes were significantly down-regulated in all individuals. Mucins constitute the mucus layer which serves as the primary barrier separating epithelial cells and host tissues from the commensal microbiota^35^. Although commensal microorganisms are generally non-pathogenic, studies suggest that in the absence of dietary glycans, the proportion of mucin-degrading bacteria may increase^36^. However, ongoing degradation of the outer mucus layer by anaerobic bacteria indicates that the mucus must be continually replaced by the epithelium^35^, potentially elevating the risks to intestinal integrity. Our results showed that one of the key beneficial effects of prebiotics: by providing nutrients (prebiotics) to microbiota, their consumption of mucin is minimized. Additionally, all six oligosaccharides reduced the activity of enzymes such as N-acetylhexosaminidase which are involved in the degradation of host glycosaminoglycans^37^. In short, the six different oligosaccharides similarly guided the microbiota toward a functional state characterized by enhanced carbohydrate metabolism and a general reduction in the degradation of host-derived components and proteins.

We consistently observed significant upregulation of substrate-specific initial-cleaving CAZymes in response to different oligosaccharide structures. This widespread induction suggests that all three microbiomes actively recognized and responded to the provided substrates, initiating primary hydrolytic steps to release monosaccharides. Although similar expression patterns of these initial CAZymes were observed across individuals, donor-specific differences were also apparent. For instance, three fructan-degrading enzymes were detected following KES and RAF treatments. However, fructan β-fructosidase (CBM66+GH32/GH32) was present in F1 and F3, while F2 exclusively expressed β-fructofuranosidase. Likewise, six distinct CAZymes involved in xylo-oligosaccharide (XYL3) degradation were upregulated in F2, compared to only one in F1 and F3, respectively. These observations suggest that while key substrate-specific CAZymes are commonly activated, the diversity and composition of CAZyme responses vary considerably across individuals. Our OTF results from MetaX further revealed that these initial CAZymes were often produced by different microbial taxa in each person. Even when the same bacterial genera were present across microbiomes, their functional responses to a given oligosaccharide could differ. This highlights the phenomenon of functional redundancy, wherein multiple microbial taxa can perform similar biochemical roles, but their activation depends on the ecological context of the community. Moreover, although all individuals expressed similar classes of primary CAZymes in response to the same oligosaccharide, the downstream metabolic processing of the resulting breakdown products diverged between individuals. In other words, the same substrate may be funneled into different metabolic pathways depending on the host microbiota.

Our results revealed that the apparent functional redundancy in CAZyme responses may be shaped by the baseline composition of the microbiota. We observed that taxa with higher initial relative abundance were more likely to upregulate specific enzymes following oligosaccharide treatment, and this appeared to mediate subsequent increases in their abundance. For instance, F2 exhibited a higher baseline abundance of *Bifidobacterium* compared to F1 and F3. Correspondingly, *Bifidobacterium* in F2 actively responded to various oligosaccharide treatments and contributed to the production of relevant CAZymes, whereas in F1 and F3, *Bifidobacterium* was not the primary contributor of those enzymes. Similarly, *Prevotella* had a notably higher baseline abundance in F1, and it was only in this individual that *Prevotella* expressed GH51 in response to ARA treatment. These patterns suggest that taxa with higher initial abundance are more likely to dominate enzyme production following substrate exposure. Notably, Bifidobacterium displayed a sharp increase in relative abundance in F2 after treatments with ARA, ISO, KES, and RAF. Our data suggest that this expansion may be linked to the efficiency and specificity of the enzymes produced. For example, in response to ARA, *Bifidobacterium* expressed endo-1,5-α-L-arabinanase (EC 3.2.1.99), rather than α-L-arabinofuranosidase (EC 3.2.1.55). In response to ISO, it predominantly expressed oligo-1,6-glucosidase rather than α-glucosidase (EC 3.2.1.20), and in response to KES and RAF, it favored β-fructofuranosidase (EC 3.2.1.26) over fructan β-fructosidase (EC 3.2.1.80). These enzymes, uniquely expressed by Bifidobacterium in F2, are more efficient in processing the respective substrates^26–28^, potentially explaining the observed bloom in this genus.

These findings suggest that while functional redundancy exists across individuals, the specific taxa responsible for enzyme production and their responses to oligosaccharides are shaped by the underlying microbial community composition. As a result, shifts in dominant taxa and their substrate utilization strategies can occur rapidly (within 18 hours), and in our study, these changes were not determined solely by the structure of the oligosaccharides. This means different microbial communities may contribute to substantial variation in the metabolic efficiency of oligosaccharide interventions across individuals. This highlights the importance of considering individual microbiota features to optimize prebiotic responses. Personalized strategies that align oligosaccharide types with host-specific microbiome features may enhance the effectiveness of dietary modulation.

Additionally, our results revealed that certain species, such as *Bifidobacterium pseudocatenulatum*, *Bacteroides uniformis*, and *Agathobacter rectalis* can respond to multiple oligosaccharide treatments. These results suggest that these species play a crucial role in oligosaccharide metabolism due to their flexible enzyme production in response to different treatments. This adaptability indicates that these species may be key contributors to carbohydrate breakdown and utilization in the gut, supporting microbiome function under varying dietary conditions. Previous studies using model human microbiomes have demonstrated that Bacteroidota members harbor a greater number and diversity of CAZyme families, suggesting a broader substrate spectrum^14^. However, our metaproteomic results offer a complementary perspective by showing that enzyme expression patterns can vary across conditions and individuals, underscoring the value of protein-level data in capturing the context-dependent functional responses of gut microbes. These results further highlight the unique value of metaproteomics in uncovering functional dynamics that cannot fully be detected through taxonomic or genomic profiles alone. Microbial taxa can alter their metabolic output independently of changes in abundance, and metaproteomic data enables the separation of taxonomic changes from functional changes, offering a more accurate view of microbial responses to dietary interventions (Figure 5e).

When we further examined certain exceptions such as the lack of a marked increase in relative Bifidobacterium abundance in F2 following XYL3 treatment, in contrast to its strong response under most other treatments, we found that *Bifidobacterium* may have limited capacity to utilize secondary metabolites derived from this substrate. Additionally, MetaX-based analysis revealed that species which were not initial CAZymes producers played a role in the downstream utilization of these secondary metabolites such as xylose. These findings underscore cross-feeding^30^ and metabolic interdependence within the microbiome, wherein distinct taxa contribute to different stages of carbohydrate utilization. This layered organization likely enhances the community’s overall metabolic flexibility and efficiency in response to complex substrates.

Of course, our results also revealed some interesting patterns. For example, GH32 was upregulated under all treatments in F1. This may suggest a broader inducibility of this enzyme under carbohydrate-rich conditions. One possible explanation is that F1’s microbiota was already primed for GH32 expression due to habitual dietary intake of fructans or similar substrates, leading to a baseline presence of this CAZyme. The further increase upon exposure to other oligosaccharides may reflect cross-induction effects or co-regulation mechanisms within carbohydrate degradation pathways, whereby general carbohydrate availability stimulates a broader set of glycoside hydrolases regardless of strict substrate specificity.

Overall, this study provides a detailed and individualized view of how the human gut microbiome functionally responds to structurally distinct oligosaccharides. By integrating metaproteomic profiling with substrate-specific enzyme analysis, we captured context-dependent microbial activities that may not be apparent through taxonomic or genomic profiling alone. The identification of both common and host-specific response patterns, including enzyme-level adaptations and potential cross-feeding interactions, highlights the added value of functional metaproteomics in advancing our understanding of microbiome-diet interactions. However, several limitations should be acknowledged. First, the sample size was limited to three young adult females, so future work involving larger and more demographically diverse cohorts will be valuable for capturing broader patterns of inter-individual variation and identifying population-level trends. While the use of an ex vivo fermentation model provided a controlled and reproducible environment to characterize microbial responses, extending this approach to in vivo settings would enable the integration of host–microbiota interactions and capture long-term ecological dynamics more comprehensively. Moreover, longer cultivation periods could help reveal slower or secondary microbial adaptations that may be critical for understanding sustained prebiotic effects. Future research incorporating larger, age-diverse cohorts and in vivo validation of host outcomes will be essential to extend these findings toward clinical and nutritional applications.

## Material and Methods

### Stool samples collection and microbial culturing with oligosaccharides

Stool samples from three healthy women were used in this study: Female 1 (F1, 33 years old), Female 2 (F2, 33 years old), and Female 3 (F3, 30 years old), which were approved by the Ottawa Health Science Network Research Ethics Board at the Ottawa Hospital (20160585-01 H). Stool samples were collected and live microbiome stored according to our validated protocol^17,38^ Briefly, stool was collected at home into anaerobic stool stabilization buffer (10% (v/v) glycerol (Sigma-Aldrich, USA), 1X phosphate-buffered saline (PBS: pH 7.6) (Fisher Scientific, USA), 1% (w/v) L-Cysteine (Sigma-Aldrich, USA) using our stool collection kit and transported on ice for processing and storage. Upon arrival, the stool was diluted and homogenized to 20% (w/v) with anaerobic stool stabilization buffer using glass beads and vortexing, followed by 5 min centrifuge at 100g, and 100μm vacuum filtering before storing at −80°C.

In summary, fecal samples from three healthy adult female donors were incubated ex vivo with six structurally distinct oligosaccharides: *1,5-alpha-L-Arabinotetraose* (CAS no. 190852-24-5), *1-kestose* (CAS no. 470-69-9), *D-(+)-raffinose pentahydrate* (CAS no. 17629-30-0), *Isomaltotriose* (CAS no. 3371-50-4), *D-Cellotriose* (CAS no. 33404-34-1), *1,4-β-D-xylotriose* (CAS no. 47592-59-6) (BIOSYNTH, USA). Stock compounds were dissolved in 1×PBS (pH 7.6) to 100 mg/ml. The RapidAIM (rapid analyses of an individual microbiome) culturing and protein extraction protocol followed that of Li et al^17^. A total of 100μL of stool, 100μL of chemical solution (100μL 1×PBS used as vehicle control), and 900μL of culture medium (Supplementary Data S2) were added to each well of a 2 mL 96-well plate with triplicates. The plates were placed on a shaker and incubated at 500 rpm and 37 °C for 18 hours under anaerobic conditions. After incubation, samples were centrifuged and pellets resuspended in 1X PBS (pH 7.6) to collect the microbial pellets^17^.

### Peptides preparation and LC-MS/MS loading

Proteins from microbial pellets were extracted according to Li et al^17^: 150 μL lysis buffer (8 M urea and 4% (w/v) SDS (sodium dodecyl sulfate) (Sigma-Aldrich, USA) in 100 mM Tris (Tris(hydroxymethyl)aminomethane)-HCl (pH 8.0)) (Milipore, USA) was added to samples and sonicated by ultra-sonicator (QSonica, cat. no. Q700MPXC) at 10 kHz in an alternating 10-second on/off pulse mode for a total of 20 min (corresponding to 10 min of effective sonication). A recirculating chiller maintained at 8 °C was employed throughout to prevent sample overheating. The SP3 (single-pot, solid-phase-enhanced sample preparation)^39^ protocol was used to process proteins for tryptic digestion. Briefly, 20μg of protein (quantified using the DC Assay kit) was first reduced with 10 mM dithiothreitol (DTT; Sigma-Aldrich, USA) at 56 °C for 30min. Subsequently, 20 mM iodoacetamide (IAA; Sigma-Aldrich, USA) was added for alkylation at room temperature for 30 min in the dark. Following protein reduction and alkylation, 10μL of magnetic beads (a 1:1 mixture of Sera-Mag Speed Beads A [Thermo Scientific; Cat. No. 09-981-121] and B [Thermo Scientific; Cat. No. 09-981-123], both Magnetic Carboxylate Modified, at a final concentration of 20 μg/μL) were added to the sample. The mixture was incubated on a shaker at room temperature for 20 min at 600 rpm to facilitate protein binding prior to magnetic separation and washing. To remove salts, the samples were washed once with 80% acetonitrile (Fisher Scientific, USA), followed by two washes with 100% ACN, each for 5 minutes at room temperature. Proteins were tryptic digested on beads with addition 40uL 50mM ABC (ammonium bicarbonate) (Sigma-Aldrich, USA), containing 0.4μg trypsin (Sigma-Aldrich, USA) (1:50 protein:trypsin) and 0.1% (w/v) FA (formic acid) (Sigma-Aldrich, USA) and incubaion at 37 ℃ for 18 hours. After bead removal by centrifugation, peptides were analyzed using an UltiMate 3000 RSLCnano system coupled to an Orbitrap Exploris 480 mass spectrometer (Thermo Fisher Scientific, USA). Peptides were loaded onto a reverse-phase analytical column and separated using a 60-minute gradient from 5% to 35% ACN in 0.1% FA at a flow rate of 300 μL/min. The mass spectrometer was operated in data-dependent acquisition mode with a full MS scan range of 350–1200 m/z, followed by MS/MS for the top 15 most intense ions with dynamic exclusion enabled.

### Peptides searching, annotation and analysis

Raw data obtained from mass spectrometry were subsequently processed using Metalab-MAG^40^ with default parameters, employing the pFind open search workflow^41^. The data were searched against the UHGP (unified human gastrointestinal protein) of MGnify database^42^, which includes 4,744 genomes from the human gut microbiome, to identify peptides and proteins, as well as to annotate taxonomic and functional information. MetaX^25^ was used for connecting taxa and function within annotated peptides. ENZYME - Enzyme nomenclature database^43^ and dbCAN^44^ were used for enzymes and CAZymes analysis, Kyoto encyclopedia of genes and genomes (KEGG) was used for functional module analysis^45^. Deseq2^46^ was employed for differential expression analysis: proteins were considered significantly differentially expressed if they exhibited a log_2_|fold change|≥ 1 and an FDR-adjusted *p-value* (*q-value*) < 0.05. WGCNA^29^ was conducted using the WGCNA R package with default settings to identify co-expressed protein modules. Module–trait correlations were used to detect those associated with oligosaccharide treatments. Pairwise comparisons between treatment groups were performed using unpaired two-sided t-tests (*n* = 3 per group). *p* < 0.05 was considered significant. Differences among groups were analyzed using one-way ANOVA followed by Tukey’s test for multiple comparisons. *p* < 0.05 (adjusted) was considered significant. Statistical tests and graphing were performed using GraphPad Prism (version 9.5.1). To assess the consistency between taxonomic and functional responses, we calculated a consistency score defined as: Consistency score = log2(FC_function_ / FC_taxon_), where FC_taxon_ is the fold change of a given taxon (treatment vs. control), and FC_function_ is the fold change of its associated functional proteins. An absolute consistency score greater than 2 was considered indicative of substantial difference in the magnitude of change between taxonomic and functional levels.

## Data Availability

The mass spectrometry proteomics data have been deposited to the ProteomeXchange Consortium *via* the PRIDE^47^ partner repository with the dataset identifier: PXD064733.

## Acknowledgments

This study received funding from the NSERC-CREATE Technologies for Microbiome Science and Engineering (TECHNOMISE) program under grant number CREATE-497995-2017, which was awarded to D.F. The authors gratefully acknowledge the assistance of ChatGPT, Grammarly, DeepL and DeepSeek in enhancing the English language presentation of this manuscript. The instrument illustration in workflow was generated with the assistance of ChatGPT.

## Author contributions

A.Z. designed and performed the experiments, processed and analyzed the data, prepared the figures, and wrote the original draft. Q.W. processed the data and contributed to data analysis and figure visualization. J.M. designed and supervised the study, contributed to the conceptualization and methodology, reviewed and revised the manuscript. Z.N. was responsible for mass spectrometry operation and reviewed the manuscript. H.Q. contributed to the Methods section and reviewed the manuscript. A.D. assisted with sample provision and experimental procedures during the RapidAIM process and reviewed the manuscript. D.F. designed and supervised the project, contributed to its conceptual development and methodology, secured funding, reviewed and revised the manuscript.

## Supplemental data

This article contains supplemental data (Supplementary Data S1: Significant KEGG modules Supplementary Data S2: Culture media reagent list).

